# Structural insights into the galanin receptors signaling

**DOI:** 10.1101/2022.01.17.476689

**Authors:** Wentong Jiang, Sanduo Zheng

## Abstract

Galanin is a biologically active neuropeptide, and functions through three distinct G protein-coupled receptors (GPCRs), namely GALR1, GALR2 and GLAR3. GALR signaling plays important roles in regulating various physiological processes such as energy metabolism, neuropathic pain, epileptic activity, and sleep homeostasis. GALR1 and GALR3 signal through the G_i/o_ pathway, whereas GALR2 signals mainly through the G_q/11_ pathway. However, the molecular basis for galanin recognition and G protein selectivity of GALRs remains poorly understood. Here, we report the cryoelectron microscopy structures of the GALR1-G_o_ and the GALR2-G_q_ complexes bound to the endogenous ligand galanin or spexin. The galanin peptide mainly adopts an alpha helical structure, which binds at the extracellular vestibule of the receptors, nearly parallel to the membrane plane without penetrating deeply into the receptor core. Structural analysis combined with functional studies reveals important structural determinants for the G protein selectivity of GALRs as well as other class A GPCRs. In addition, we show that the zinc ion is a negative allosteric regulator of GALR1 but not GALR2. Our studies provide insight into the mechanisms of G protein selectivity of GPCRs and highlight potential novel function of the neuromodulator zinc ion as a modulator of GPCR signaling in the central nervous system.

**Significance Statement:** Galanin exerts various physiological functions through galanin receptors, including antinociceptive activity, depression and sleep. Here, we reveal a distinct binding site and binding pose of galanin peptide in galanin receptors from that of the published structures of peptide-bound GPCRs. Moreover, our work show that the neuromodulator zinc ion negatively modulates galanin signaling in the central nervous system, and further advances our understanding of mechanisms of G protein selectivity of GPCRs. These unique features of galanin receptors can be exploited for rational design of subtype selective ligands for treatments of neurological disorders.

## Introduction

Galanin is a 29 or 30-amino-acid peptide that was isolated from pig intestine in 1983 (1). Through its wide distribution in the nervous system and the endocrine system, galanin is involved in a variety of physiological functions, including regulation of hormones and neurotransmitters release, antinociceptive activity, depression and sleep/wake homeostasis (2, 3). The endogenous action of galanin is mediated through activation of three distinct receptor subtypes (GALR1-3), which belong to the class A of G protein-coupled receptors (GPCRs) family (4, 5).

GALR subtypes vary in their downstream signaling pathways and the tissue distribution. GALR1 and GLAR3 mainly couple to the inhibitory Gα_i/o_ pathway, leading to the inhibition of the adenylyl cyclase activity and the decrease of the intracellular adenosine 3’,5’-cyclic monophosphate (cAMP) level. By contrast, GALR2 mainly couples to the stimulatory pathway of G_q/11_, inducing the formation of inositol triphosphate (IP3), which in turn increases the cytosolic Ca^2+^ level (3) **(Fig. 1*A*).** While GALR1 is particularly enriched in the nervous system, GALR2 and GALR3 are broadly distributed in brain as well as peripheral tissues. GALRs activation via overexpression or administration of galanin in the nervous system of animals suppresses seizure development and neuropathic pain behavior, and show anxiolytic and antidepressant effect (6–10). A missense mutation in galanin peptide was identified as a cause of temporal lobe epilepsy (TLE) (11). Moreover, galanin expression is upregulated in the injured neurons, and galanin has been shown to play a role in neuroprotection and neuronal regeneration (12, 13). Accumulating evidence indicate that GALRs signaling is a key regulator of both sleep time and sleep/awake homeostasis in model organisms such as zebrafish and mouse (14, 15). Therefore, GALRs are potential therapeutic targets for the treatment of pain, epilepsy, depression, neuron injury and sleep disorders.

**Fig. 1.**
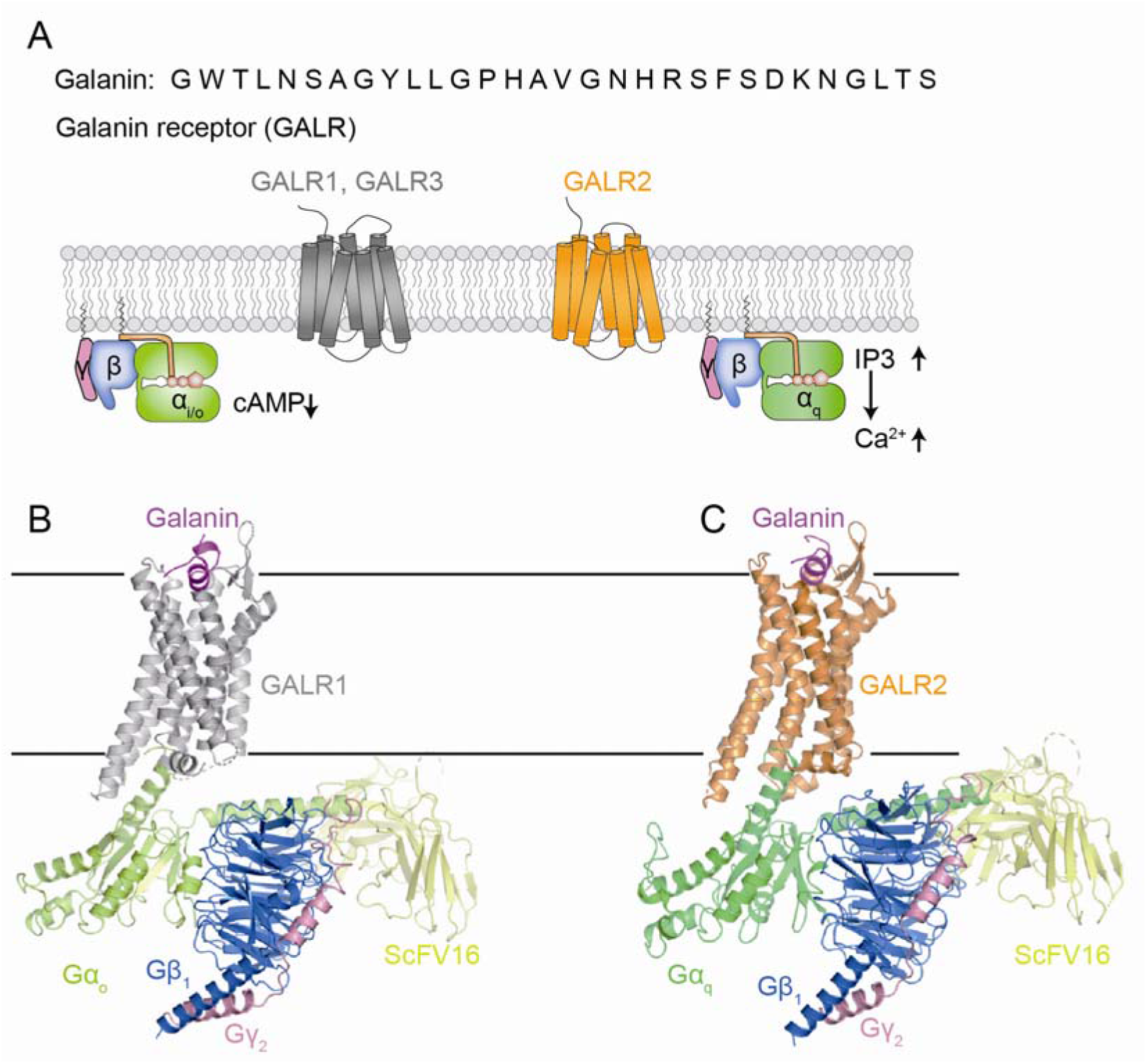
Overall structures of galanin-bound GALR1-miniGo-scFv16 and GALR2-miniGq-scFv16 complexes. (A) Schematic representation of GALR receptors signaling. GALR1 and GALR3 primarily couple to Gi/o, while GALR2 mainly signals through Gq. (B) Cryo-EM structures of GALR1-miniGo-scFv16. (C) Cryo-EM structures of GALR2-miniGq-scFv16.

In addition to galanin, the endogenous galanin-like peptide (GALP) and spexin have been shown to activate GALRs (16, 17). Although these peptides share high sequence similarity, they show distinct receptor binding preference. In contrast to galanin that interacts with all three receptor subtypes, spexin specifically activate GALR2 and GALR3. Homology modeling and site directed mutagenesis studies revealed the essential residues of galanin involved in the receptor binding and activation, and the potential galanin binding site of GALRs (18, 19). However, the molecular details of galanin binding and the peptide selectivity of GALRs remain poorly defined at the molecular level. Moreover, little is known about the molecular basis of G protein coupling specificty of GALRs. To gain insight into the molecular basis of ligand recognition and ligand selectivity of GALRs and extend our understanding of G protein selectivity, we sought to determine the cryoelectron microscopy (cryo-EM) structures of GALR1 and GARL2 in complex with G_o_ and G_q_ heterotrimer respectively.

## Results

### Structure determination

To obtain stable GPCR-G protein complexes, we used engineered thermostable mini-G proteins, which only contain the GTPase domain of Gα but still bind to Gβγ heterodimer and recapitulate the pharmacological and structural changes in GPCRs induced by the full-length Gα proteins (20). Moreover, the N-terminal residues of αN in mini-Gαo and mini-Gaq were replaced by the equivalent residues of Gαi to acquire the ability to bind the antibody fragment scFv16 that stabilizes the nucleotide-free GPCR-G protein complex (21). Furthermore, we introduced a linker that contains a 3C protease cleavage site, between the C-terminus of the receptor and the N-terminus of the mini-Gα to create a GPCR-G fusion protein. The GALR1-mini-Gαo or GALR2-mini-Gαq fusion protein was transiently expressed in Expi293 cell, and was assembled with purified Gβ1γ2 and scFv16 in the presence of galanin **(*SI Appendix*, Fig. S1).** The resulting GALR1-mini-Go complexes were co-eluted and mono-disperse with or without 3C protease treatment from the size exclusion chromatography, indicating that GALR1 forms a stable complex with mini-Go **(*SI Appendix*, Figs. S1A-1C**). The peak fraction corresponding to the complexes were concentrated and subjected to cryo-EM single particle analysis. 2D class average analysis showed that the GALR1-mini-G fusion protein complex gives more orientations than the GALR1-mini-G complex without a linker between the receptor and min-Gα (***SI Appendix*, Figs. S1D and S1E**). Combination of the two datasets enables us to obtain a final cryo-EM map of the GALR1-mini-Go complex at a global nominal resolution of 3.3 Å (***SI Appendix*, Fig. S2 and Table S1**). The structure of the galanin- and spexin-bound GALR2-minGq fusion complex was determined to a nominal resolution of 3.3 Å and 3.5 Å, respectively **(*SI Appendix*, Fig. S3 and Table S1)**. The high quality EM map allowed us to unambiguously assign side chains of the galanin peptide 1-17 and the most amino acids of the receptors except the extreme terminal residues and some intracellular loops because of their high flexibility **(Fig. 1 *B* and C)**. The overall structure of the GALR1-Go complex resembles that of the GALR2-Gq complex, with root-mean-square deviation values of 0.886 Å for the Cα atoms of the receptors and 0.604 Å for the Cα atoms of the G proteins.

### Comparison of galanin binding pockets of GALR1 and GALR2

The existence of bulky aromatic amino acids and the high quality EM density map allowed us to unambiguously assign side chains of galanin **(Fig. 2*A*).** The N-terminal portion of galanin (residue 1-15) was well resolved due to its direct contact with the receptors, which is consistent with previous studies showing that the binding affinity of N-terminal region of galanin (1–16) for the receptors is comparable to the full-length galanin (22, 23). Moreover, the N-terminal region (1–16) but not the remaining part is highly conserved in GALP and spexin peptides **(*SI Appendix*, Fig. S4A),** both of which are able to activate the receptors, further supporting our structural observation. Galanin mainly forms an alpha helical structure when bound to the receptor as well as in solution itself (24, 25). It occupies at the extracellular vestibule of GALRs that is equivalent to the binding site of a positive allosteric agonist LY2119620 in M2R (26) **(*SI Appendix*, Fig. S4B)**. It lays on top of the receptor, nearly parallel to the membrane plane and distant from the toggle switch W^6.48^, the conformational change of which is essential for the receptor activation. By contrast, most neuropeptide agonists of class A GPCRs such as endothelin, orexin and opioid peptides binds nearly perpendicular to the membrane plane with one end buried in the helical cavity and the other end interacting with the extracellular loops, and these peptides penetrate in proximity to the toggle switch (27–29) (***SI Appendix*, Fig. S4B**). Galanin contacts all seven TM helices as well as extracellular loops ECL2 and ECL3, burying a surface area of 866 Å^2^, which accounts for the high affinity binding of galanin for GALRs in the sub-nanomolar range (30). GALR1 and GALR2 use overlapping but distinct set of residues to contact galanin, mostly via hydrophobic and hydrogen bond interactions **(Fig. 2).** The first N-terminal residue of G1 lies between TM1 and TM7, and is closer to TM1 of GALR1 than that of GALR2 **(Fig. 2*F*)**, which may explain that removal of G1 in galanin reduced its binding affinity for GALR1 but not GALR2 (24). W2 is sandwiched between L277^ECL3^ and F282^7.32^ of GALR1 and makes additional hydrogen bond with S281^7.31^ **(Fig. 2*C*).** Therefore, mutation of W2 in galanin or F282^7.32^ results in significant loss of binding for the receptors (18, 31). F282^7.32^A mutation in GALR1 almost abolished galanin potency **(Fig. 2*G*),** and F271^7.32^A mutation in GALR2 reduced galanin potency by nearly 100-fold **(Fig. 2*H*).** A7E mutation that was identified as a cause of TLE disease likely causes a clash with nearby hydrophobic residues, accounting for reduced binding affinity for GALRs (11) **(Fig. 2*C*).** Y9 penetrates into the receptor core, about 10 Å above the toggle switch (***SI Appendix*, Fig. S4B**), and is hydrogen-bonded by Q92^2.61^ in GALR1 or Q82^2.61^ in GALR2 **(Figs. 2 C and *E*).** Mutation of Q92^2.61^ or Q82^2.61^ to alanine reduced galanin potency by almost 100-fold **(Figs. 2 *G* and *H*)**. Our structural observation is also consistent with previous studies showing that W2 and Y9 are vital for galanin binding to the receptors (31). However, because of distinct residues of ECL2 and ECL3 involved in binding galanin, the conformations of these regions are different between GALR1 and GALR2 **(Fig. 2*F*).** For instance, V274 in the ECL3 of GALR1 engages hydrophobic interaction with L11 **(Fig. 2*C*),** while the equivalent residue in GALR2, Q263 rotates away from galanin due to its longer side chain and hydrophilic nature, resulting in the conformational change of ECL3 **(Fig. 2*F*)**. As a result, V274G mutation reduced agonist potency by about 370-fold, while Q263A mutation showed little effect **(Figs. 2 *G* and *H*).** ECL2 forms an antiparallel β-sheet, which is a characteristic of peptide receptors. It covers galanin as a lid-like structure and forms extensive hydrophobic interactions with L4, P13 and V16. The residues in ECL2 of GALR1 involved in binding galanin have bulkier aromatic side chains than that in GALR2 **(Fig. 2*F*)**. Mutations of the equivalent residues W188^ECL2^ and H176^ECL2^ in GALR1 and GALR2 respectively had distinct effect on galanin potency **(Fig. 2 *G* and *H*),** suggesting that ECL2 in GALR1 and GALR2 differently contribute to galanin binding. An endogenous peptide spexin has A7M and G8L mutations in galanin and specifically activates GALR2 and GALR3 **(*SI Appendix*, Fig. S4A)**. Structure of the spexin-bound GALR2-Gq complex reveals that L8 in spexin likely clashes with the bulkier residue W188^ECL2^ in GALR1 **(*SI Appendix*, Fig. S4C)**, accounting for the specific binding of spexin for GALR2 and GALR3 (16). Owing to these conformational differences, R184^5.35^ in GALR2 but not K197^5.35^ in GALR1 makes hydrogen bonds with the backbone of galanin **(Fig. 2*F*)**. As a result, R184^5.35^ A mutation reduces galanin potency by about 10-fold, while K197^5.35^ A mutation shows little effect **(Figs. 2 *G* and *H*).** Taken together, these results suggest that mechanisms of galanin recognition by GALR1 and GALR2 are not identical, which allows the development of selective ligands targeting a specific subtype.

**Fig. 2.**
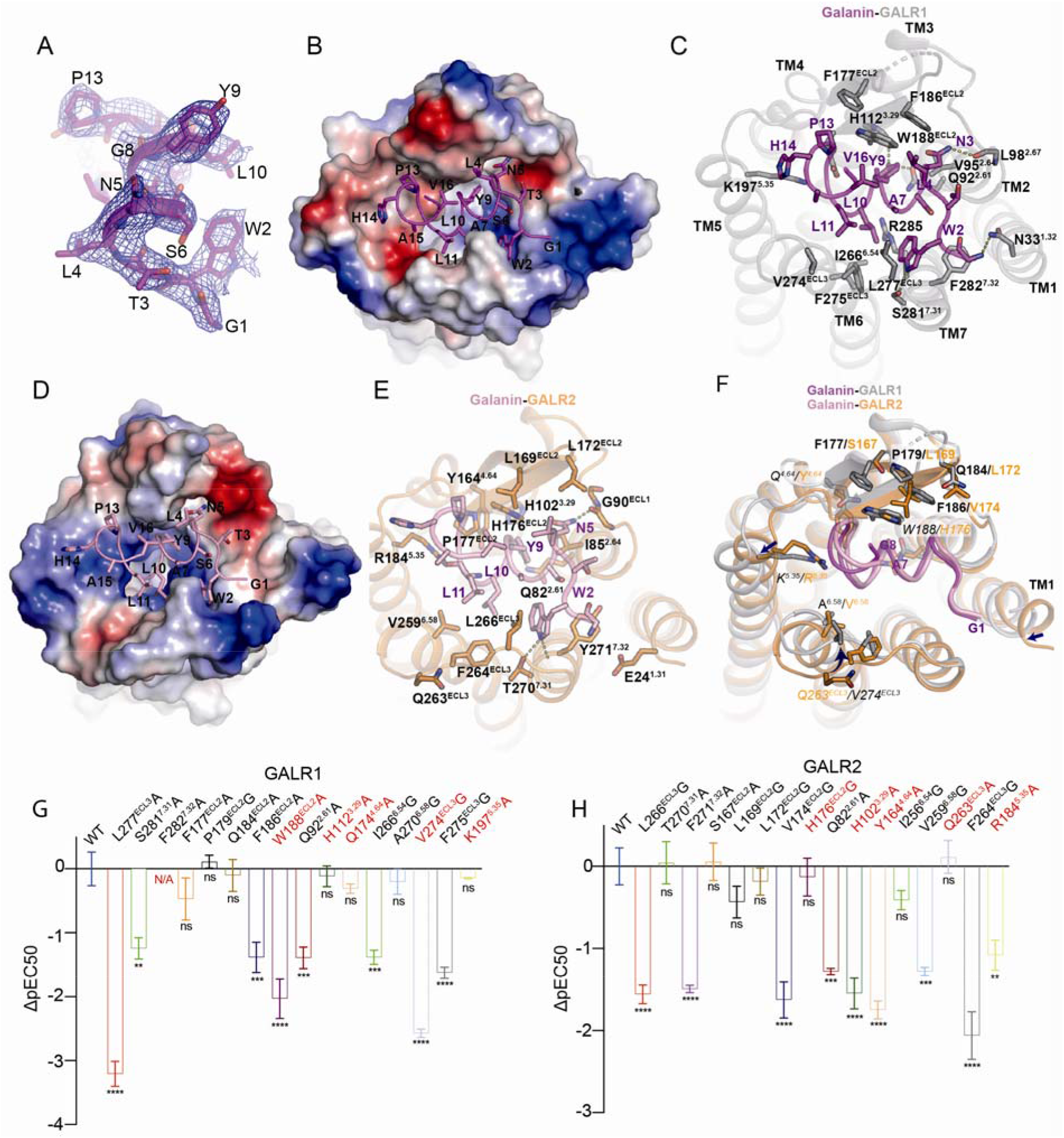
Mechanisms of galanin recognition by GALR1 and GALR2. (A) EM density map for galanin from the structure of the GALR2-Gq complex. (B) Electrostatic potential surface of GALR1 and ribbon representation of galanin (magenta) viewed from the extracellular side. Colors from red to blue represent negatively to positively charged regions. (C) Detailed interaction between GALR1 and galanin. (D) Electrostatic potential of the GALR2-galanin interface is distinct from that of the GALR1-galanin interface. (E) Detailed interaction between GALR2 and galanin. (F) Structural superposition of the GALR1-galanin and the GALR2-galanin complex. The equivalent residues in GALR1 and GALR2 that play distinct roles in galanin binding are shown. Arrows indicate the conformational changes. (G) and (H) The effects of mutations in GALR1 and GALR2 on galanin potency as measured by the cAMP inhibition assay and the IP1 accumulation assay respectively. The equivalent residues that play distinct roles in GALR signaling are colored red. Data represent mean ± SEM of triplicate measurements in three independent experiments. Significance was analyzed using one-way ANOVA, ****P<0.0001, ***P<0.001, **P<0.01.

### Activation mechanisms of GALR1 and GALR2

The inactive structures of GALR1 and GALR2 predicted by Alphafold may represent ligand-free structure, in which TM helices loosely pack against each other to allow the access of galanin (32) **(Fig. 3*A*).** Upon galanin binding, the orthosteric site undergoes significant conformational change, as indicated by the inward displacement of the extracellular portions of TM2 and TM6 and the outward shift of the TM1 and TM7 **(Figs. 3 *A* and *B*).** These conformation changes account for the outward motion of TM6 and inward motion of TM7 in the intracellular side **(Fig. 3*C*)**. The conformational changes of the toggle switch W^6.48^ and P^5.50^I/V^3.40^F^6.44^ motif are common features of the class A GPCR activation. In contrast to the most class A GPCRs, where the orthosteric sites are in close proximity to the toggle switch, galanin binding site is distant from it. Hydrophobic interactions between F275 in ECL3 of GALR1 and, L10 and L11 in galanin result in the downward shift of F275, which propagates to the downward movement of W260^6.48^ via I266^6.54^ and H263^6.51^ **(Fig. 3*B*)**. The downward shift of W^6.48^ is associated with the conformational change of the P^5.50^I/V^3.40^F^6.44^ motif, which allosterically disrupts the conserved ionic lock between R133^3.50^ and D132^3.49^, leading to the outward displacement of TM6 **(Fig. 3*C*)**. Inward displacement of TM7 in the intracellular side is observed in the active state of many other class A GPCRs, as indicated by the conformational change of NPXXY motif, in which Y303^7.53^ forms a water-meditated hydrogen bond with Y220^5.58^. The inward displacement of TM7 is coupled by the outward shift of R285^7.35^ that arises from its interaction with galanin. Although the key residues involved in receptor activation are conserved between GALR1 and GALR2, their conformations vary significantly **(Fig. 3*D*).** This is because the hydrophobic interaction between V274 in ECL3 and L11 in galanin exists in GALR1, while this interaction is absent in GALR2 due to the substitution of V274 in Q263, which leads to the upward shift of F264^ECL3^ as well as the toggle switch W^6.48^ and PIF motif in GALR2, compared to the equivalent residues in GALR1 **(Fig. 3*D*).** To further support the important role of F275^ECL3^/F264^ECL3^ in GALR receptors activation, mutation of F275 in GALR1 or F264 in GALR2 reduced galanin potency by almost 100-fold **(Figs. 2 *G* and *H*).** The conformation differences of residues involved in receptor activation contribute to the structural variation in the cytoplasmic pocket of GALR1 and GALR2 and may play a role in G protein selectivity.

**Fig. 3.**
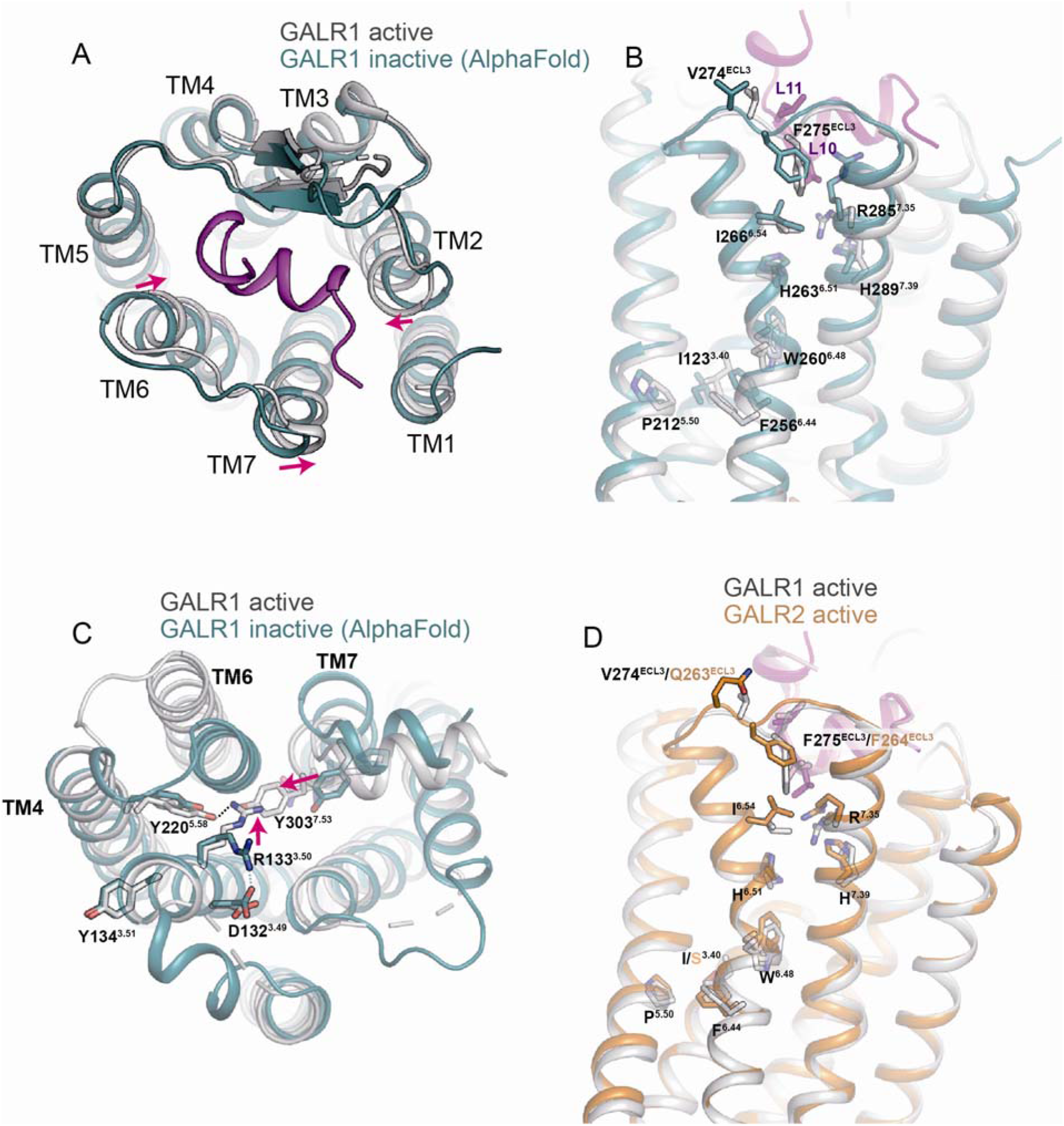
Mechanisms of GALR1 and GALR2 activation. (A) Structural overlay of GALR1 in the active and inactive state (predicted by AlphaFold). (B) Conformational changes of the P^5.50^I^3.40^F^6.44^ motif upon receptor activation. (C) Conformational changes of residues in the cytoplasmic pocket including the D^3.49^R^3.50^Y^3.51^ and the NPY motif upon receptor activation. (D) Structural overlay of the active state of GALR1 and GALR2.

### Zn^2+^ is a negative allosteric modulator (NAM) of GALR1

Previous studies have reported that Zn^2+^ can inhibit galanin binding to the receptors (18). To further investigate the functional role of zinc ion in galanin receptors signaling, we used the NanoBiT complementation-based assay to assess the effect of Zn^2+^ on activation of receptors by galanin in living cells. Mini-G proteins were used in the NanoBiT assay throughout this study, since they preserve appropriate coupling specificity, and can be recruited to the active GPCRs without further dissociation, which increases the signal-to-noise ratio in this assay. As expected, Zn^2+^ diminished the effect of 1 μM galanin on GALR1 activation in a concentration dependent manner with an IC50 value of 47.2 μM **(Fig. 4*A*).** The diminished effect of Zn^2+^ is saturable, or has a “ceiling” level. In contrast, the diminished effect was observed in GALR2 when the concentration of Zn^2+^ reached millimolar range that is above the physiological concentration, indicating that Zn^2+^ had little effect on galanin-induced GALR2 activation. Our structures show that galanin receptors are enriched with histidine residues that may coordinate Zn^2+^ underneath the orthosteric binding pocket **(Fig. 4*B*).** Comparison of primary sequences of GALR1 and GALR2 from different species revealed that H267^6.55^ but not nearby histidine residues in GALR1 is mutated to Isoleucine in GALR2 **(*SI Appendix,* Fig. S5A)**. As expected, the zinc effect was significantly abrogated in GALR1 when H267^6.55^ was mutated, while H112^3.29^A, H263^6.51^F or H264^6.52^F mutation had little influence **(Fig. 4*A* and *SI Appendix*, Fig. S5B)**. All these mutants of GALR1 can be activated by galanin, although the potency and efficacy of galanin for these mutants vary **(*SI Appendix*, Fig. S5C)**. We further tested the effect of Zn^2+^ on the concentration response curve of galanin. Zn^2+^ produced the concentration-dependent and saturable rightward shifts in the potency of galanin, and decreased the galanin maximum response as well **(Fig. 4*C*).** By contrast, Zn^2+^ had little effect on the galanin concentration-response curve of H267A mutant of GALR1 **(Fig. 4*D*)**. H267 is located in the TM6 right below the galanin binding site. The extracellular part of TM6 near H267 moves inwards upon galanin binding, which leads to the receptor activation. As a result, when coordinated by H267 and other nearby residues, Zn^2+^ likely rigidifies the extracellular part of TM6 and restricts its conformational change, attenuating galanin-induced receptor activation. The exact coordination pattern of Zn^2+^ awaits further investigation. Nevertheless, these results indicate that Zn^2+^ is a NAM of GALR1.

**Fig. 4.**
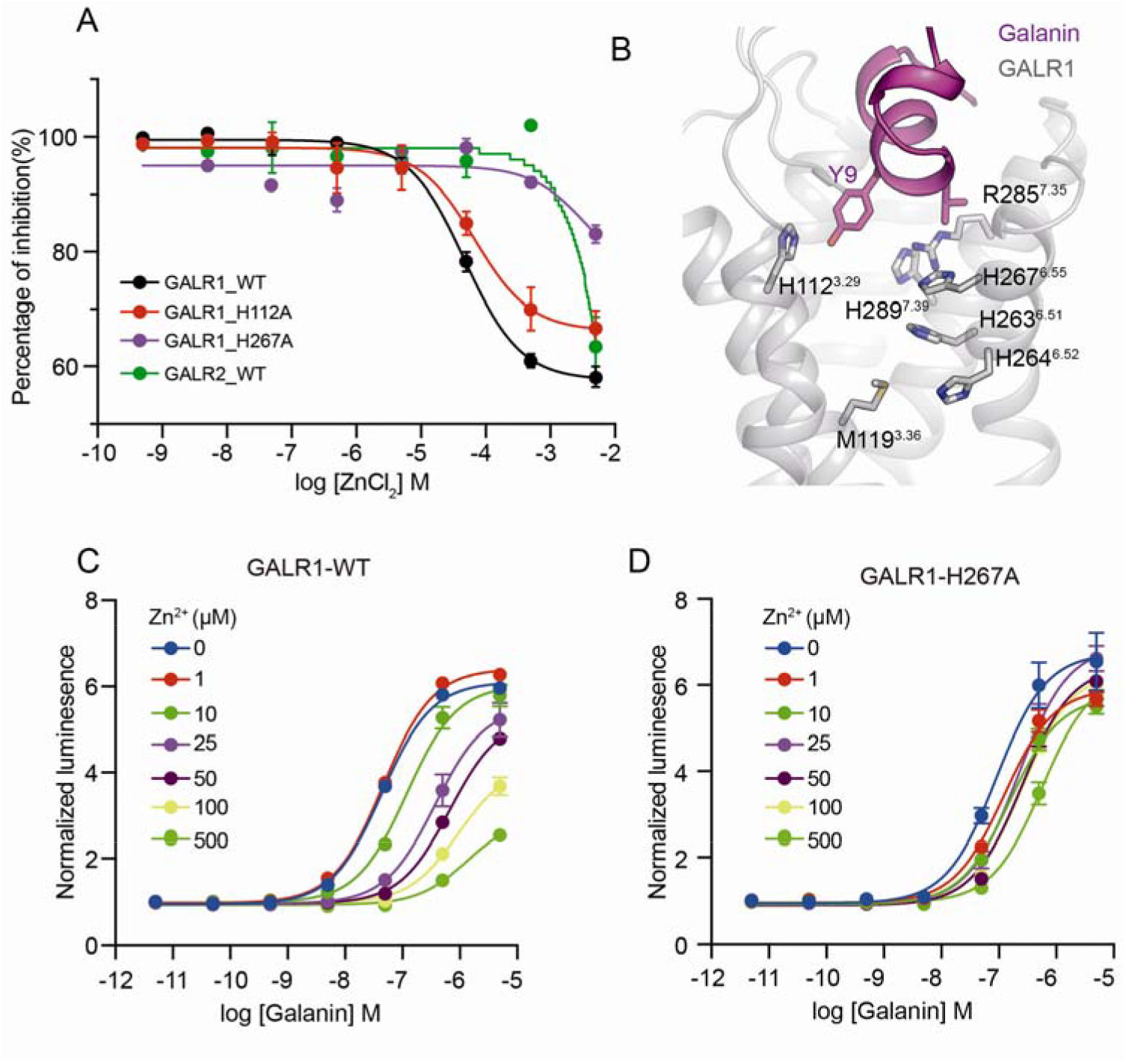
Zinc is a NAM of GALR1. (A) The effect of increasing concentration of zinc on receptor activation induced by 1 μM galanin, as evaluated by the NanoBiT assay, where the small fragment, and the large fragment are fused to the C-terminus of GALR1 and the N-terminus of mini-Go, respectively. The luminescence signals are normalized as percentages of the initial response of GALR1 to galanin without zinc treatment. (B) Histidine residues are enriched underneath the galanin binding pocket of GALR1. (C) and (D) The actions of increasing concentration of zinc on the galanin dose-response curve of WT (C) and H267A mutant (D) of GALR1 measured by the NanoBiT assay. The luminescence signals are normalized to the vehicle treatment as fold change.

### Structural determinants of Gi/o and Gq/11 selectivity

A notable difference between structures of the GALR1-Go complex and the GALR2-Gq complex is the relative orientation of Go and Gq to the receptors **(Figs. 5 *A* and *B*).** When aligning the receptors, the α5 of Gαo is rotated around the “wavy hook” of α5 by about 14° toward TM5, compared with Gαq. This orientation difference was also observed in the structures of M1 and M2 muscarinic receptors (M1R and M2R) bound to G11 and Go respectively (33). In addition, because of the different interaction interface of the receptor and G protein, the flexibility of intracellular loops (ICL) in GALR1 and GALR2 differs **(Figs. 5*A*).** For instance, ICL1 is ordered in GALR2 owing to the hydrogen bond interaction between D312 in Gβ and the main chain carbonyl group of G53 in ICL1, whereas it is flexible in GALR1 due to the absence of this interaction **(*SI Appendix*, Fig. S6A)**. The ICL2 of most Gi/o-coupled receptors forms an alpha helical structure, where hydrophobic residues at position 34.51 engage weak hydrophobic interactions with the hydrophobic pocket of Gαi/o formed by V34 from the αN-β1 loop, L195 from the β2-β3 loop and I343 and F336 from α5 **(*SI Appendix*, Fig. S6B)**. However, the ICL2 of GALR1 is disordered because of the substitution of the hydrophobic residue L1313^4.51^ in arginine and the absence of hydrophobic interaction between the ICL2 of GALR1 and Gi **(Figs. 5 *C* and *G*).** By contrast, L131^34.51^ in the ICL2 of GALR2 is buried deep in the hydrophobic pocket of Gαq formed by L34 from the αN-β1 loop, V79 from the β2-β3 loop, and F228, K232 and I235 from α5, and engages strong hydrophobic interactions. In addition, P130^34.50^ at the junction of ICL1 and TM3 is stabilized through hydrophobic interactions with I235 and K232 in α5 of Gαq **(Fig. 5*D*).** Substitution of L131^34.51^ or P130^34.50^ in GALR2 in the equivalent residues in GALR1 impaired the ability of GALR2 to couple Gq **(*SI Appendix*, Fig. S7A)**, accounting for the inability of GALR1 to couple Gq. However, substitutions of S140^34.50^, R141^34.51^ and S149^4.49^ in ICL2 of GALR1 with the equivalent residues in GALR2 have little effect on coupling efficiency between GALR1 and Gαi **(*SI Appendix*, Fig. S7B)**, suggesting that ICL2 in GALR1 is not involved in Gi coupling. Remarkably, GALR1 acquires the ability to couple Gq, as indicated by the NanoBiT assay as well as the IP1 assay **(Fig. 5*H* and *SI Appendix*, Fig. S7C)**, when residues in ICL2 of GALR1 are substituted with that of GALR2, further supporting the important role of ICL2 in Gq coupling.

**Fig. 5.**
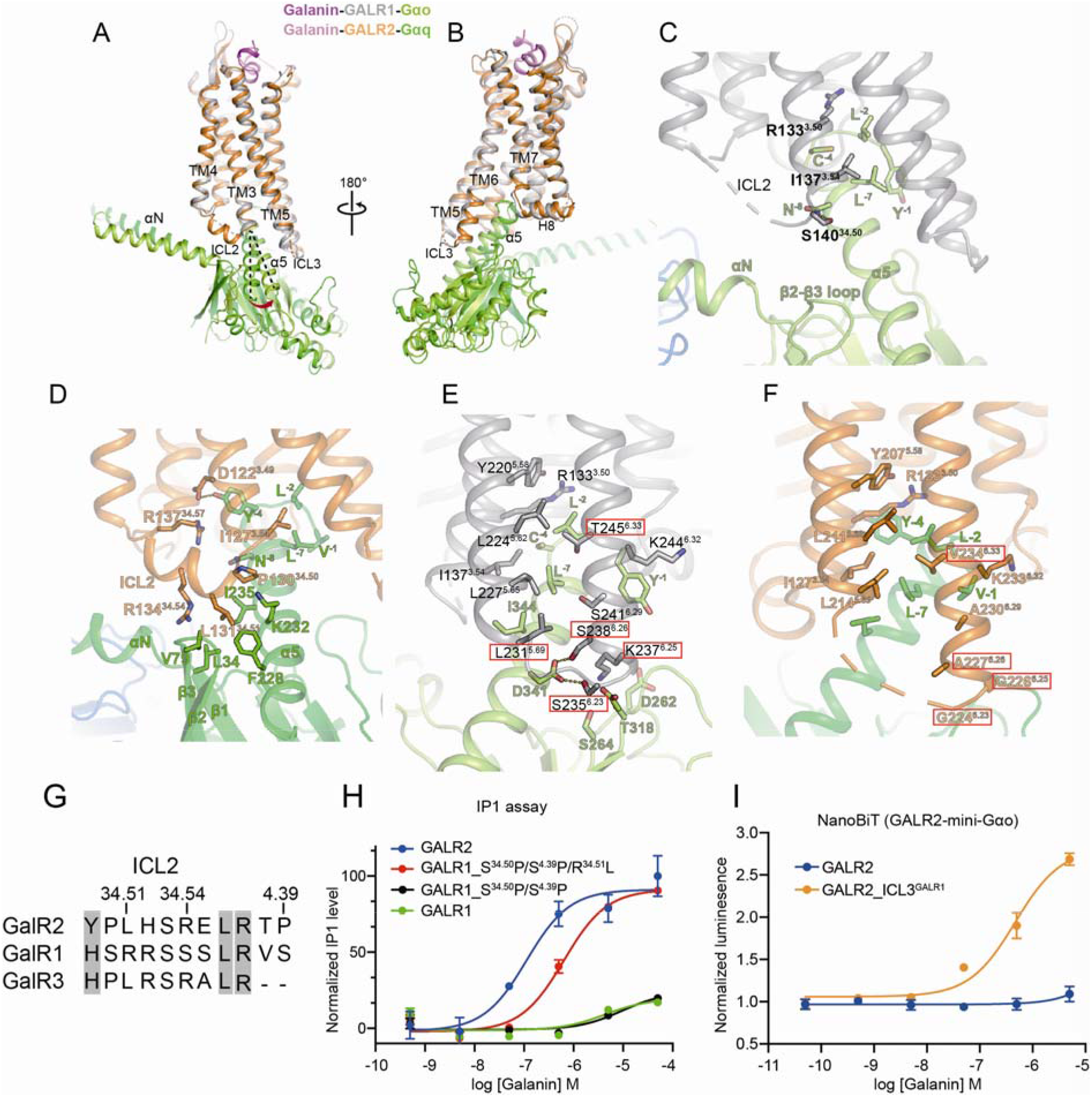
Mechanisms of Go and Gq selectivity in the GALR receptor family. (A) and (B) Structural superposition of GALR1-Gαo and GALR2-Gαq in two opposite views. Receptors are aligned. (C) Interaction details between ICL2 of GALR1 and Gαo. (D) Interaction details between ICL2 of GALR2 and Gαq. (E) Interaction details between TM5 and TM6 of GALR1 and Gαo. (F) Interaction details between TM5 and TM6 of GALR2 and Gαq. (G) Sequence alignment of ICL2 from GALR receptor family. (H) The IP1 accumulation assay evaluating the effects of ICL2 substitutions in GALR1 on GALR1-Gq coupling. All mutants are expressed at similar levels as WT. (I) Substitution of ICL3 in GALR2 with that in GALR1 increases coupling efficiency of GALR2 and Go.

To understand the structural mechanism of the inability of GALR2 to couple Gi, we compared the interaction details between the GALR1-Gi and the GALR2-Gq complexes and mainly focused on residues of GALR1 involved in Go coupling that are not conserved in GALR2. R133^3.50^, I137^3.54^, L224^5.62^, L227^5.65^, L231^5.69^, and T245^6.33^ in TM3, TM5 and TM6 of GALR1 engage extensive hydrophobic interactions with the extreme C-terminal part of α5 in Go **(Fig. 5*E*).** Most of these residues are conserved in GALR2 **(Fig. 5*F*).** Mutations of conserved residues in GALR1 and GALR2 impair the recruitment of Gi and Gq, respectively (***SI Appendix*, Figs. S7D-7G**). Nevertheless, a notable difference between GALR1 and GARL2 are in ICL3. S238^ICL3^ and S235^ICL3^ in ICL3 of GALR1 make hydrogen bonds with D341 in Gαo, and K237^ICL3^ engages electrostatic interactions with residues in the GTPase domain of Gαo. All of three residues are mutated in GALR2, leading to loss of these interactions **(Fig. 5 *E* and *F*)**. Mutations of S235^ICL3^ and K237^ICL3^ in GALR1 have modest effect on galanin potency, but dramatically reduced the maximum responses (***SI Appendix*, Figs. S7D and S7E**). Remarkably, GALR2 acquired the ability to bind Go, when ICL3 (214-225) of GALR2 including the three residues were replaced by the equivalent residues in GALR1 **(Fig. 5*I*).** Taken together, our data suggest that ICL2 in GALR2 and ICL3 in GALR1 are critical for determining the Gq and Go selectivity, respectively.

### Structural determinants of Gs and Gq selectivity

Although it has been shown that interactions between the hydrophobic residue at position 34.51 of ICL2 and the hydrophobic pocket of Gα are essential for the efficient coupling of Gq and Gs (34), it remains unclear how Gq and Gs are selectively recognized. Comparison of structures of D1 dopamine receptor (D1R)-Gs and GALR2-Gq revealed key structural elements in the receptors that determine Gq and Gs selectivity. In the GALR2-Gq complex, the conformation of ICL2 is stabilized by salt bridge interactions between R^34.57^ (M^34.57^ in D1R) and D^3.49^ of the DRY motif as well as a hydrogen bond between R^34.57^ and Y(−4) in Gαq, while ICL2 of D1R is stabilized by a hydrogen bond between Y^34.53^ (S^34.53^ in GALR2) and D^3.49^, and a potential water-meditated hydrogen bond between Y^3.49^ and Y(−4) in Gas **(Fig. 6*A*).** Notably, Y^34.53^M/V^34^.^57^ are prevalent in Gs-coupled receptors, while R^34.57^ is enriched in Gq-couple receptors **(Fig. 6*D*).** Mutations of YM in D1R and RS in GALR2 significantly reduced the potency of dopamine and galanin, respectively **(Figs. 6 *E* and *F*)**. Moreover, N(−3) (−1 indicates the last residue of Gα) in Gαq is inserted into a hydrophobic pocket formed by N^2.40^, F^8.50^ and other nearby residues, whereas E(−3) flips outside this pocket, probably due to its longer side chain and negative-charge nature. Remarkably, when E(−3) in Gas but not the nearby residues L(−1) and Q(−5) was substituted with the equivalent residues in Gαq, the coupling efficiency between GALR2 and Gs was significantly increased **(Fig. 6*G*).** These different interaction modes of GALR2-Gq and D1R-Gs account for the movement of α5 in Gs toward TM6 and the outward movement of TM6 in D1R, compared to that in GALR2 **(Fig. 6*B*),** explaining that most Gs-coupled receptors display a larger TM6 movement in the active state than Gq-coupled receptors. As a result of these conformational changes, Gs is closer to TM5 than Gq, highlighting the important role of TM5 in determining Gs selectivity. Indeed, TM5 in most Gs-coupled receptors have a C-terminal helical extension, and previous studies have shown that the A/V^5.65^ Q^5.68^ϕ^5.69^ (ϕ represents hydrophobic residues) motif in TM5 is prevalent in receptors that exclusively couple to Gs and is critical for Gs coupling in D1R (35) (Companion paper). Residues at position 5.65 in Gs-coupled receptors prefer hydrophobic residues with small side chains such as alanine and valine because of its close distance from the hydrophobic pocket formed by L(−1), L(−2), and L(−7) in Gas **(Fig. 6*C*)**. Mutation of A^5.65^ in leucine would cause a clash with this pocket, and impaired the Gs coupling (Companion paper). In contrast, leucine is dominant at position 5.65 in Gq-coupled receptors, due to its long distance from the hydrophobic pocket formed by V(−1), L(−2), and L(−7) in Gαq **(Fig. 6*C*)**. L^5.65^A mutation in GALR2 weakens the interaction with this hydrophobic pocket and thus significantly decreased galanin potency **(Fig. 6*F*).** However, it is noteworthy that Gs- and Gq-coupled receptors show sequence preference at some positions of ICL2 and TM5, but also accommodate various residues at these positions **(Fig. 6*D*),** partly because of diverse receptor-G protein interfaces and promiscuous coupling of some GPCRs.

**Fig. 6.**
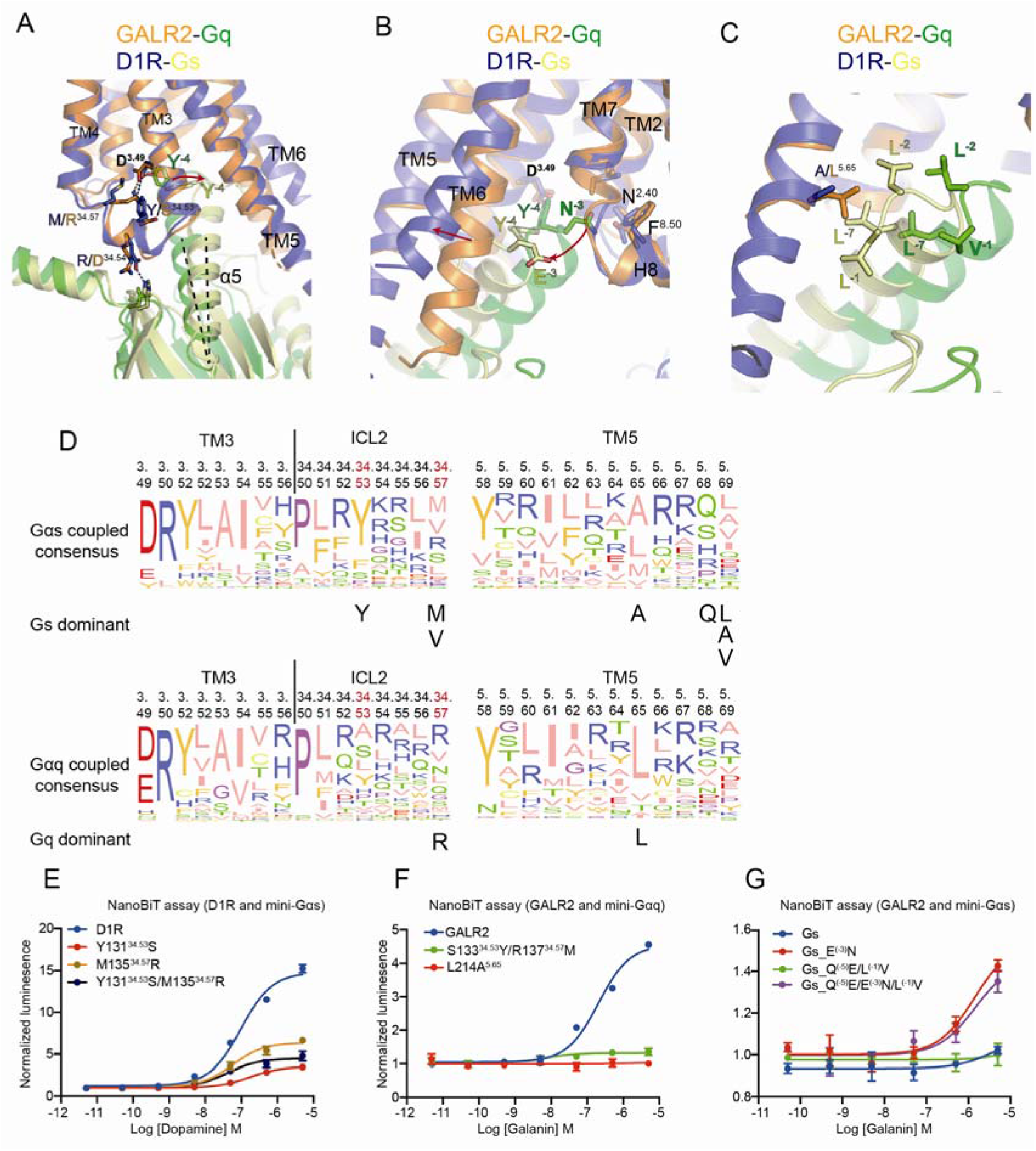
Important structural features in class A GPCRs that determining Gs and Gq selectivity. (A) Structural superposition of the GALR2-Gq and the D1R-Gs complexes. Receptors are aligned. (B) N(−3) in Gq is inserted into a hydrophobic pocket formed by TM2, TM7 and H8, while E(−3) in Gs flips outside this pocket. (C) A^5.65^ in D1R is close to the hydrophobic pocket formed by L(−1), L(−2) and L(−7) in Gs, while L^5.65^ in GALR2 is distant from that in Gq. (D) Sequence alignment of 41 class A Gs-coupled receptors (top) and 44 Gq-coupled receptors (bottom). The dominant residues are indicated below the alignments. (E) Mutations of Y^34.53^M^34.57^ in the ICL2 of D1R reduce D1R-Gs coupling efficiency. (F) Mutations of S^34.53^R^34.57^ in the ICL2 of GALR2 almost abolish GALR2 and Gq coupling. (G) The effects of mutations of the “wavy hook” in Gs on coupling efficiency of GALR2-Gαs, as evaluated by the NanoBiT assay.

## Discussion

Here, we report cryo-EM structures of the GALR1-Go and GALR2-Gq complex using the GPCR-G protein fusion strategy. The structures revealed distinct mechanisms of galanin recognition and receptor activation for GALR1 and GALR2, which contribute to structural variation in the cytoplasmic pocket of the receptors and may play an important role in determining the G protein selectivity. Moreover, we showed that Zn^2+^ is a negative allosteric modulator of GALR1 but not GALR2.

Zn^2+^, known as a neuromodulator, is widely distributed in the central nervous system (CNS), particularly enriched in the synaptic vesicles of glutamatergic neurons (36, 37). It is released to the synaptic cleft upon membrane depolarization, and modulates functions of ion channels and receptors on the pre- or post-synaptic membrane. It has been shown that zinc inhibits ionotropic glutamate AMPA and NMAR receptors, fine-tuning synaptic transmission in the brain (38, 39). GALR1 is expressed on both glutamatergic and GABAergic postsynaptic neurons. The spatial colocalization of zinc and GALR1 makes it possible for zinc to regulate the function of GALR1. Moreover, the IC50 of zinc on GALR1 activation is 47.2 μM, which is in the range of the physiological concentration of zinc (10 nM to 100 μM) (40). Previous studies have also shown that zinc regulates endogenous ligand binding at several GPCRs including β2 adrenergic receptors (β2AR) (41), melanocortin receptors (42) and platelet-activating factor receptor (43). In this study, we showed that zinc attenuated GALR1 activation by galanin possibly through restricting the conformational change of TM6 that leads to receptor activation. Further studies are required to address whether zinc modulates a large number of GPCRs in the CNS and fine-tunes GPCR signaling, as does sodium (44).

Combining published structures of the GPCR-Gi complexes, we can roughly divide the class A Gi-coupled receptors into three classes based on the interaction features between ICL2 and G proteins: (i) receptors that exclusively couple to Gi and have a charge residue at the position 34.51 of ICL2 such as GALR1 and sphingosine-1-phosphate receptors (S1PR) (45); (ii) receptors that exclusively couple to Gi and have a hydrophobic residue at the position 34.51 such as D3 dopamine receptor, M2 muscarinic receptor (M2R) and μ opioid receptor (28, 33, 46); (iii) receptors that promiscuously couple to Gi and have a large hydrophobic residue at position 34.51 such as the neurotensin receptor 1 (NTSR1), β2AR, and the cholecystokinin A receptor (47, 48). In the first class, when bound to the receptor, ICL2 is disordered, or forms a random coil structure. Since there is no hydrophobic interaction between 34.51 and Gαi, ICL2 in receptors of this class plays a distinct role in determining Gi coupling efficiency (45) (***SI Appendix*, Fig. S8A**). In GALR1, ICL2 is not involved in Gi coupling, whereas in S1PR, ICL2 is involved in hydrophilic interactions with Gi, and is important for Gi coupling; In the second class, ICL2 forms an alpha helical structure, and the residue 34.51 of ICL2 is located outside and distant from the hydrophobic pocket of Gαi formed by residues from the αN-β1 loop, the β2-β3 loop and α5, and engages weak hydrophobic interactions (***SI Appendix*, Figs. S8B and S8C**). Mutation of this residue had little effect on the Gi coupling or GDP release from Gi/o (49, 50); In the third class, similar to Gs- and Gq-coupled receptors, the residue 34.51 is located close to the middle of the hydrophobic pocket of Gαi and engages strong hydrophobic interaction (***SI Appendix,* Figs. S8E and S8F**). In addition, some receptors in this class such as NTSR1 have the other conformation, where the residue 34.51 is located outside the hydrophobic pocket (***SI Appendix,* Fig. S8D**). Previous studies have shown that F^34.51^A mutant of β2AR failed to activate Gi (50), suggesting that the hydrophobic interaction between ICL2 and Gi is very important for Gi coupling in the third class. Owing to the absence of or weak interaction between Gαi and ICL2 in receptors that exclusively couple Gi, the cytoplasmic end of TM5 and TM6, and ICL3 have strong interactions with Gαi, and are critical for determining Gi selectivity **(*SI Appendix*, Fig. S8).**

It has been recognized that the distal part of α5 in Gα plays a key role in determining G protein selectivity (51–53). We further identified a residue pair N/E(−3) in α5 of Gq/Gs that contributes to structural differences and selective interactions between the D1R-Gs and GALR2-Gq. Substitution of this residue in Gs can promote coupling of GALR2 to noncognate Gs. Moreover, we revealed several signature residues in ICL2 and TM5 that dominate in Gs- and Gq-coupled receptors. Thus, our results provide novel insights into the molecular mechanisms of G protein selectivity by class A GPCRs.

## Materials and method

### Cloning

The human GALR1 and GALR2 were cloned into pcDNA3.1(+) vector(Thermo Fisher Scientific) with an N-terminal hemagglutinin (HA) signal sequence and a FLAG epitope tag (DYKDDDDK). An engineered mini-Gαo was fused to the C-terminus of GALR1 (1-349) with three copies of 3C protease sites between them. GALR2 (1-314) was expressed as a fusion protein including two repeats of 3C protease site and a mini-Gαq sequence in the C-terminus of GALR2. ScFv16 was cloned into the pFastBac vector (Invitrogen) with an N-terminal GP64 signal sequence and a C-terminal 3C protease site, followed by an octa-histidine tag. His6-tagged Gβ1 and Gγ2 (C68S mutation) were cloned in the pFastBac Dual vector for insect cell expression.

### Protein expression and purification

The plasmid expressing GALR1-mini-Gαo or GALR1-mini-Gαq was transiently expressed into Expi293F cells (Thermo Fisher Scientific) using polyethyleneimine (PEI, Polysciences). Cells were lysed in the lysis buffer (20 mM HEPES, pH 7.4) supplemented with protease inhibitor cocktail (Roche) using a glass dounce grinder and centrifuged at 1,000 x g for 3 minutes to remove the nucleus. The membrane fraction was pelleted by centrifugation at 65,000x g, at 4 °C for 1 hour, and homogenized in the solubilization buffer containing 20 mM HEPES pH 7.4, 150 mM NaCl, 1% Lauryl Maltose Neopentyl Glycol (LMNG), 0.2% Cholesteryl Hemisuccinate (CHS) and 60 nM galanin peptide (1–30). After centrifugation to remove debris, the supernatant containing solubilized GALRs-mini-Gα was supplemented with 2 mM CaCl_2_ and loaded onto the M1 anti-FLAG antibody resin. The resin was washed with wash buffer containing 20 mM HEPES, pH 7.4, 300 mM KCl, 0.01% LMNG, 0.002% CHS, 2 mM CaCl_2_, 10 mM MgCl_2_, 2 mM ATP, 6 nM galanin and eluted with elution buffer containing 20 mM HEPES, pH 7.4, 150 mM NaCl, 0.01% LMNG, 0.002% CHS, 10 mM EDTA, 0.5 mg/ml 1x FLAG peptide and 60 nM galanin. Gβ_1_γ_2_ (C68S) and scFv16 were expressed and purified as previously described (21, 54).

### Complex assembly

Purified GALR1-miniGo or GALR2-miniGq proteins, Gβ_1_γ_2_ (C68S) and scFv16 were mixed with a molar ratio of 1:1.5:2 in 500 μl of the equilibration buffer (20 mM HEPES, pH 7.4, 150 mM NaCl, 0.01% LMNG, 0.002% CHS, 60 nM galanin, 0.5 μM TCEP) supplemented with 1 μl PNGaseF and 0.5 μl apyrase. The mixture was incubated on ice for 1 hour and further purified on a Superose 6 Increase 10/300 column pre-equilibrated with the equilibration buffer. The peak fractions containing the complex were supplemented with 60 μM galanin and concentrated to about 6 mg/ml. For assembly of the GALR1-miniGαo/Gβ1γ2 (C68S) complex with 3C protease site cleaved, similar procedures were performed as above, except that 3C protease was added before purification on a Superose 6 Increase 10/300 column. For assembly of the spexin-bound GALR2 complexes, same procedures were performed, except that galanin was replaced by spexin during the purification process.

### Cryo-EM sample preparation and data collection

300 mesh holey carbon grids (Quantifoil Au R1.2/1.3) were glow-charged, loaded into a Vitrobot MarkIV instrument chamber (Thermo Fisher Scientific) maintained at 8 °C and 100% humidity. 3.0 μl of GALRs complex samples was applied onto the grid, blotted for 3.0-4.0 s with a blotting force of 4 before plunge freezing in liquid ethane. Cryo-EM movies were collected on a Titan Krios microscope equipped with a BioQuantum GIF/K3 direct electron detector (Gatan) under accelerating voltage of 300 kV at a nominal magnification of 64,000 x. Each movie stack was collected as 32 frames, with total dose of 50 e-/Å^2^ for 2.56 s.

### Cryo-EM Data processing

All movie stacks were collected and processed with MotionCor2 for motion correction (55), with 2x binned to a pixel size of 1.087 Å. Contrast Transfer Function (CTF) estimation was performed using patch-based CTF estimation in cryoSPARC_v3 (56). All processed images were then subjected to particle picking using Blob picker in cryoSPARC, followed by particle extraction. For the galanin-bound GALR1-mini-Go complex with 3C protease sites cleaved, particles from 1,401 micrographs (Dataset A) were subjected to two rounds of 2D classification, generating 248,352 good particles. *Ab initio* reconstruction and non-uniform refinement were performed to get a reference map for GALR1. For the GALR1-mini-Go fusion protein complex, 886 micrographs (Dataset B) were collected, followed by particle picking using Blob picker and particle extraction. Particles from the two datasets were combined and subjected to two rounds of 2D classification, yielding 2,882,487 good particles. These particles were subjected to global 3D classification in RELION3.1 (57), followed by another round of 3D classification focused on the receptor. 426,045 particles from the best class were run through non-uniform refinement in cryoSPARC, resulting in a final 3.3 Å map.

For the galanin-bound GALR2-mini-Gq fusion protein complex, 1,337 micrographs were collected, and processing procedures were performed as above. In brief, two rounds of 2D classification using auto-picked particles resulted in 1,325,739 good particles, which were subjected to two rounds of 3D classification in RELION3.1 using the GALR1-Go complex map as initial model. 578,453 particles from three classes with clear secondary structure features were selected, and subjected to non-uniform refinement in cryoSPARC, resulting in a final 3.29 Å map. All 3D maps were post-processed with DeepEMhancer (58).

For the spexin-bound GALR2-mini-Gq complex, 1,139 movies were collected and processed as above. 1,015,461 good particles were selected from two rounds of 2D classification, and were subjected to heterogeneous refinement and non-uniform refinement in cryoSPARC. The final map is about 3.5 Å.

### Model building

Homology models for GALR1 and GALR2 were generated using the structure of μ opioid receptor (PDB: 4DKL) in the SWISS-MODEL server. The homology model of GALR1 and the structure of mini-Gαo/Gβγ/scFv16 (PDB: 7D77) were fitted into the EM map in Chimera (59). The structure of mini-Gαq/Gβγ/scFv16 was extracted from the published structure (PDB: 6WHA), and docked into the EM map together with the homology model of GALR2. All the models were manually built in COOT (60) and, are subjected to *real_space_refinement* in *Phenix* (*61*) using the reference structure and secondary structure restraints. The statistics for structure refinement are summarized in Table S1.

### cAMP inhibition assay

Chinese hamster ovary (CHO) cells were seeded into six-well plates and cultured overnight until cell confluence reaches ~80%. Plasmids expressing GALR1 or mutants were transfected together with the GloSensor biosensor plasmid following a Lipofectamine cell transfection procedure (Invitrogen). Transfected cells were cultured for 1 day and re-seeded into 96-well plates by 3 x 10^4^ cells per well. After 8 hours post seeding, the medium was exchanged to CO2-independent medium (Gibco) supplemented with 500 μg/ml of D-luciferin. Cells were stimulated with various concentration gradients of galanin for 5 minutes, and then treated with 1 μM forskolin. The bioluminescence signal was constantly measured for 10 minutes, and the peak signal was acquired for the inhibitory dose curve fitting and IC50 determination using GraphPad Prism 8 software. Significance analysis was performed using one-way analysis of variance method (one-way ANOVA in Prism 8).

### IP1 accumulation assay

Gαq-mediated IP1 accumulation was measured using the IP-ONE Gq HTRF Kit from Cisbio. HEK-293T cells were seeded into 6-well plates, and 2 μg of GALR2 or mutant plasmids were transfected using PEI. After 2 days post-transfection, cells were suspended, washed one time with DPBS (Gibco), resuspended into HBSS buffer (Beyotine) and seeded into 384-well plates(Greiner) with ~7000 cells per well. Transfected cells were stimulated with various concentration gradients of galanin for 1 hour and subjected to IP1 accumulation detection following the assay protocol. Inhibitory dose curve was plotted and IC50 was determined using GraphPad Prism 8 (dose-response-inhibitory, three parameters). Significance was analyzed using One-way ANOVA.

### NanoBiT assay

To monitor the interaction between G proteins and GALR1 or GALR2 upon galanin stimulation, a NanoLuc-based enzyme complementation system called NanoBiT assay (62) was used (Promega). The C-terminus of GALR1 or GALR2 was fused with the small fragment (smBiT), and the large fragment (LgBit) element was fused to the N terminus of mini-Gα proteins. HEK-293T cells were seeded into 6-well plates and transfected with 1 μg of GPCR-smBit and 1 μg of LgBit-miniGα. After 2 days post transfection, cells were suspended, washed with DPBS for one time and resuspended into the assay buffer containing HBSS supplemented with 0.01% BSA (SIGMA), 10 mM HEPES (Beyotine) and 10 μM coelenterazine-h (YEASEN). The culture was equilibrated at room temperature (RT) for 2 hours and subjected to stimulation with various concentration gradients of galanin and instant bioluminescence measurement. The bioluminescence signal was acquired in the time point when the signal went into the stationary phase, and the normalized signal (fold change) was fitted to a three-parameter sigmoidal concentration-response curve in Prism 8 software.

### Zn^2+^ inhibition assay

As zinc produced high background signal in the IP1 accumulation assay and cAMP inhibition assay, the NanoBiT assay was used to measure the effect of Zn^2+^ effect on GALRs signaling. The same constructs used in the NanoBiT assay were adopted. After 2 days post-transfection, cells were resuspended and washed twice with the assay buffer (20 mM HEPES, pH 7.3 and 150 mM NaCl), and resuspended into the assay buffer supplemented with 10 μM coelenterazine-h and seeded into 96-well plates. After 30 minutes of incubation at RT, cells were stimulated with various concentration gradients of galanin premixed with a fixed concentration of ZnCl_2_, or 1 μM of galanin pre-mixed with titrated concentration of ZnCl_2_. The bioluminescence signals in the stationary phase were acquired, and were analyzed using three-parameter dose-response-stimulatory or dose-response-inhibitory fitting methods in Prism 8 software.

## Acknowledgements

We thank Dr. Xiangyu Liu at Tsinghua University for providing the plasmid expressing Gβ1γ2. We thank staff at Shuimu BioSciences for their help with cryo-EM data collection. All EM images were collected at Shuimu BioSciences. This work was supported by Chinese Ministry of Science and Technology, Beijing Municipal Science & Technology Commission (Z201100005320012) and Tsinghua University.

## Author contributions

W.J. purified the protein complex, collected cryo-EM data, performed cryo-EM data processing and model building, performed cellular assay with input from S.Z. S.Z. and W.J. wrote the manuscript.

## Data availability

The atomic structures have been deposited at the Protein Data Bank (PDB) under the accession codes XXX. The EM maps have been deposited at the Electron Microscopy Data Bank (EMDB) under the accession numbers XXX.

