## Supplemental material for "Structural insights into the galanin receptors signaling"

Sanduo Zheng

**This PDF file includes:**

Figures S1 to S8

Tables S1

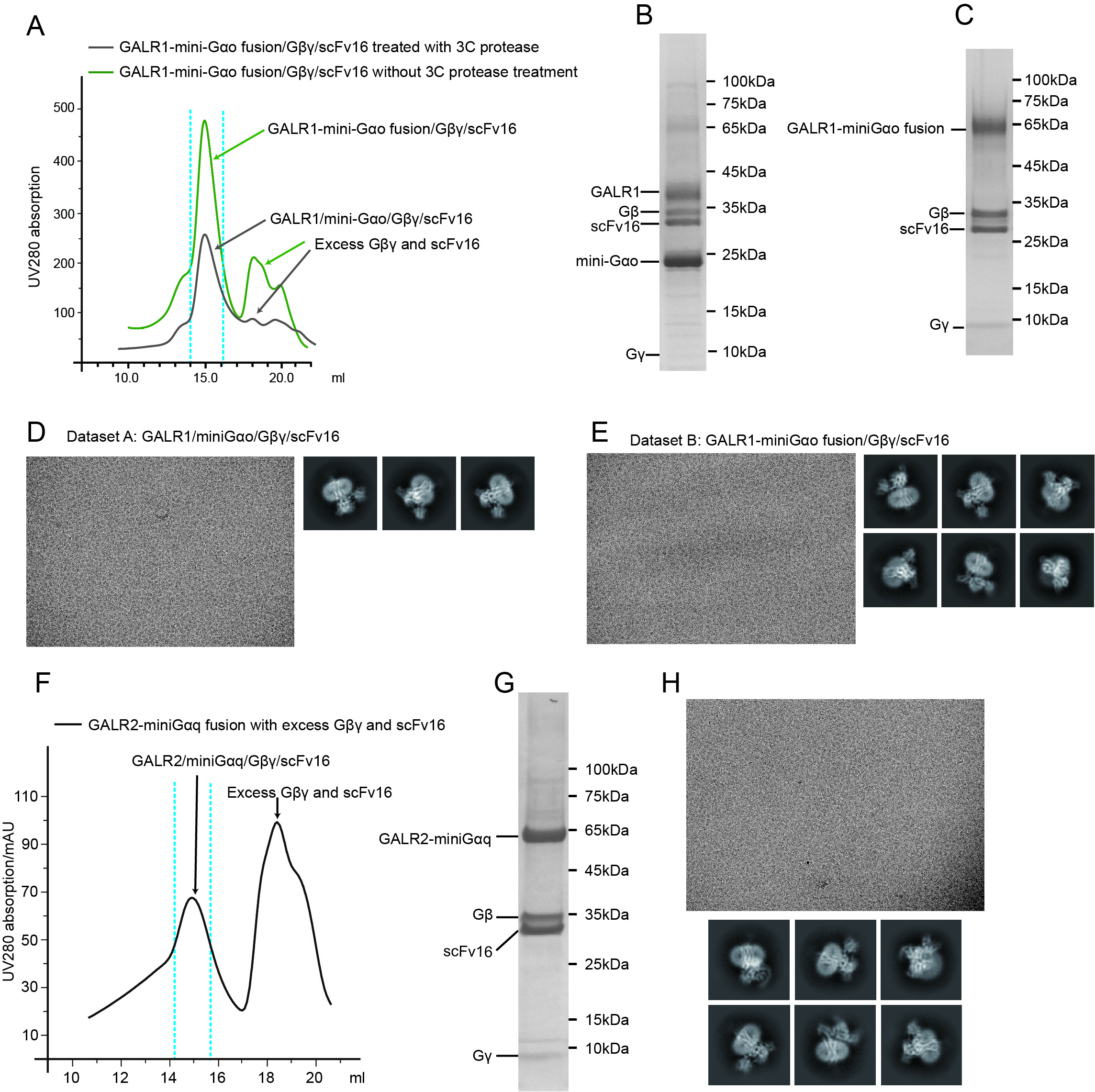

**Fig. S1.** **Protein purification and cryo-EM analysis.**

(A) Size exclusion chromatography of the complex of the GRLR1-miniGαo fusion protein bound to Gβ1γ2 and scFv16, with or without 3C protease treatment that cleaves the linker between GALR1 and mini-Gαo.

(B) and (C) SDS-PAGE of the GRLR1-mini-Go complex with (B) or without (C) 3C protease treatment.

(D) Cryo-EM images and 2D classification of the GALR1/mini-Go/ Gβ1γ2/scFv16 complex without a linker (Dataset A).

(E) Cryo-EM images and 2D classification of the complex of the GALR1-miniGo fusion protein bound to Gβ1γ2 and scFv16 (Dataset B).

(F) Size exclusion chromatography of the complex of the GRLR2-mini-Gαq fusion protein bound to Gβ1γ2 and scFv16.

(G) SDS-PAGE of the GALR2-mini-G complex.

(H) Cryo-EM images and 2D classification of the GALR2-mini-G complex.

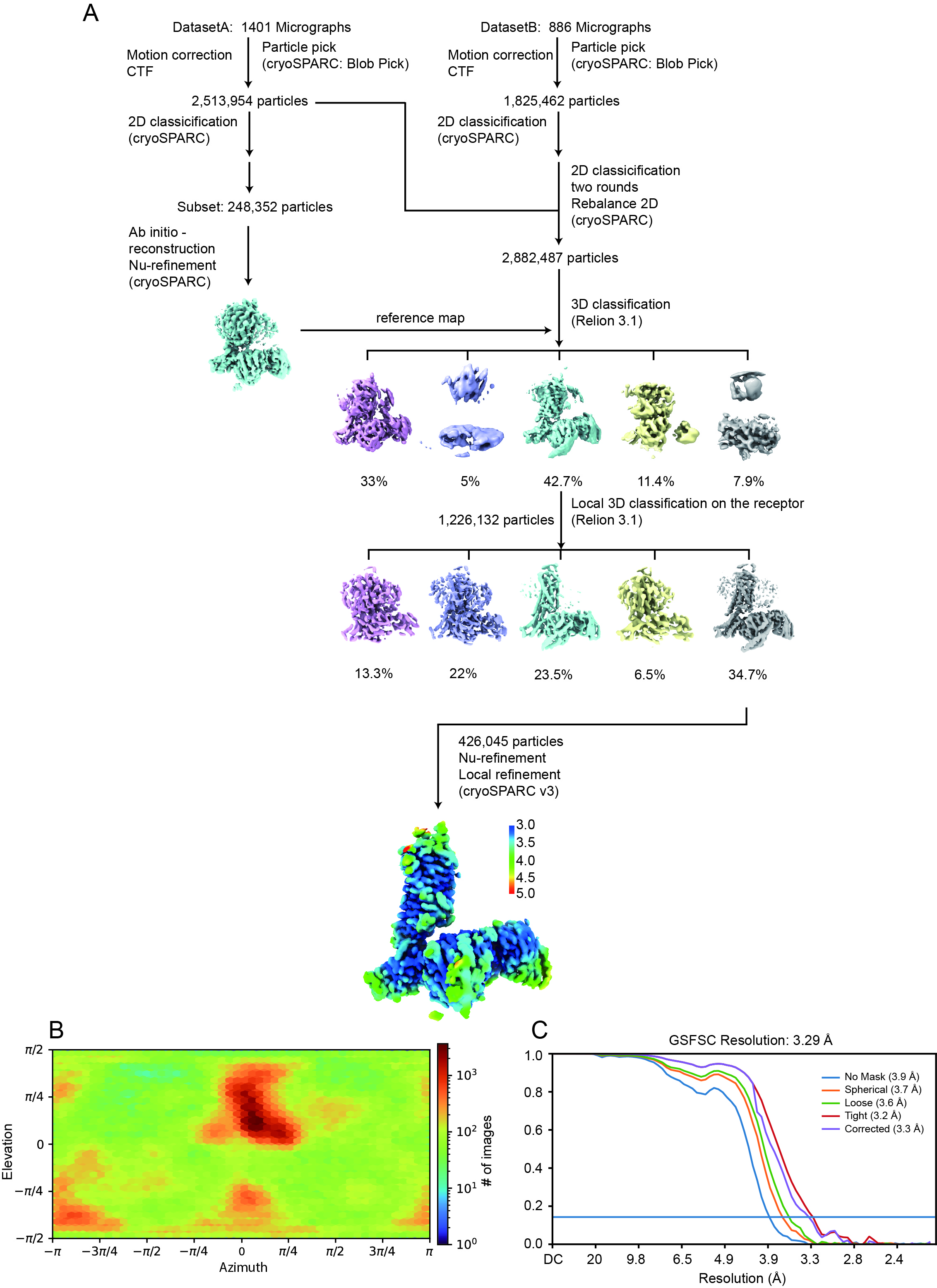

**Fig. S2. Cryo-EM data processing of the galanin-bound GALR1-mini-Go complex.** (A) Cryo-EM workflow chart.

(B) Angular distribution plot.

(C) Gold standard FSC curves.

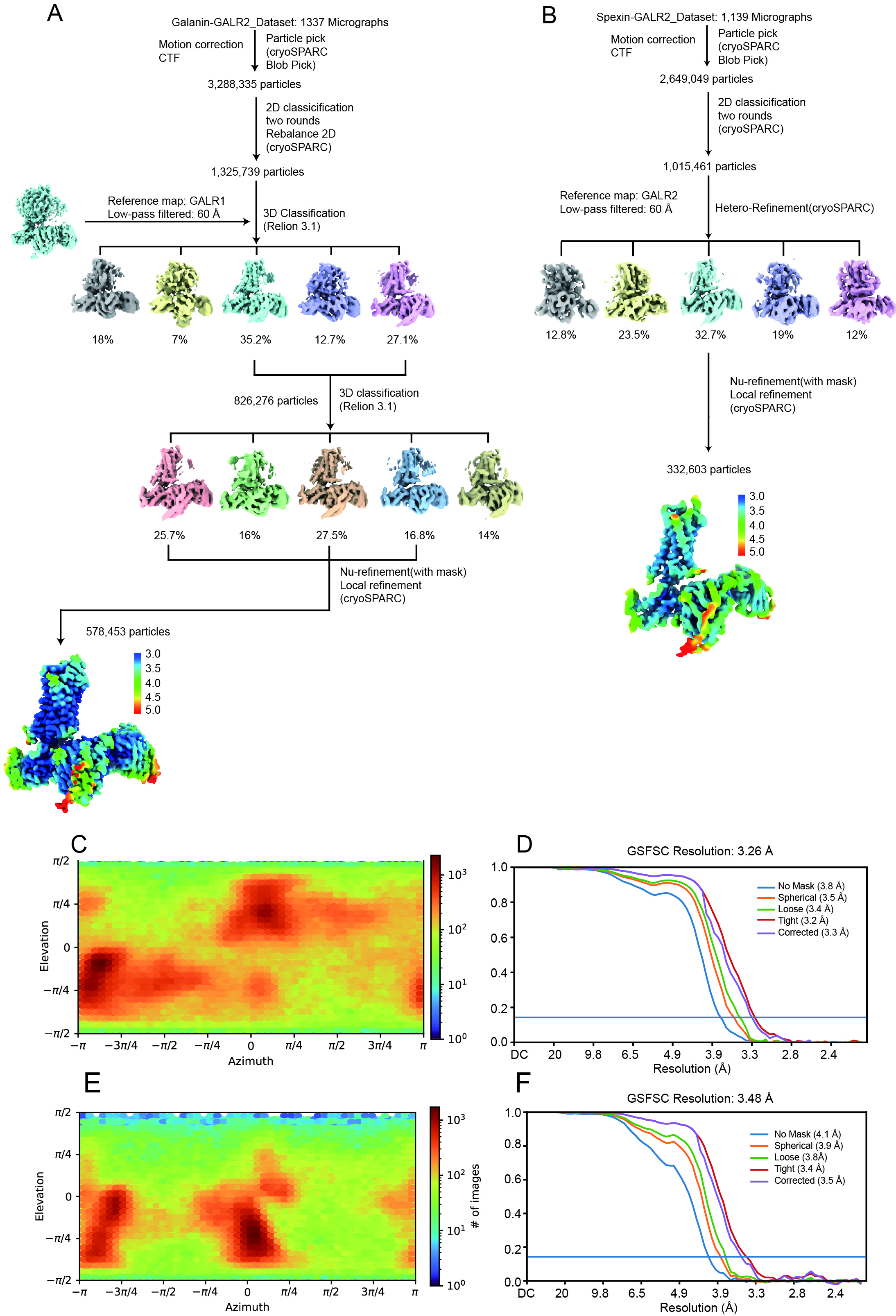

**Fig. S3.** **Cryo-EM data processing for the GALR2 complexes.**

(A) and (B) Cryo-EM workflow chart for the galanin- (A) and the spexin- (B) bound GALR2-mini-Gq complex.

(C) Angular distribution plot for the galanin-bound complex.

(D) Gold standard FSC curves for the galanin-bound complex.

(E) Angular distribution plot for the spexin-bound complex.

(F) Gold standard FSC curves for the spexin-bound complex.

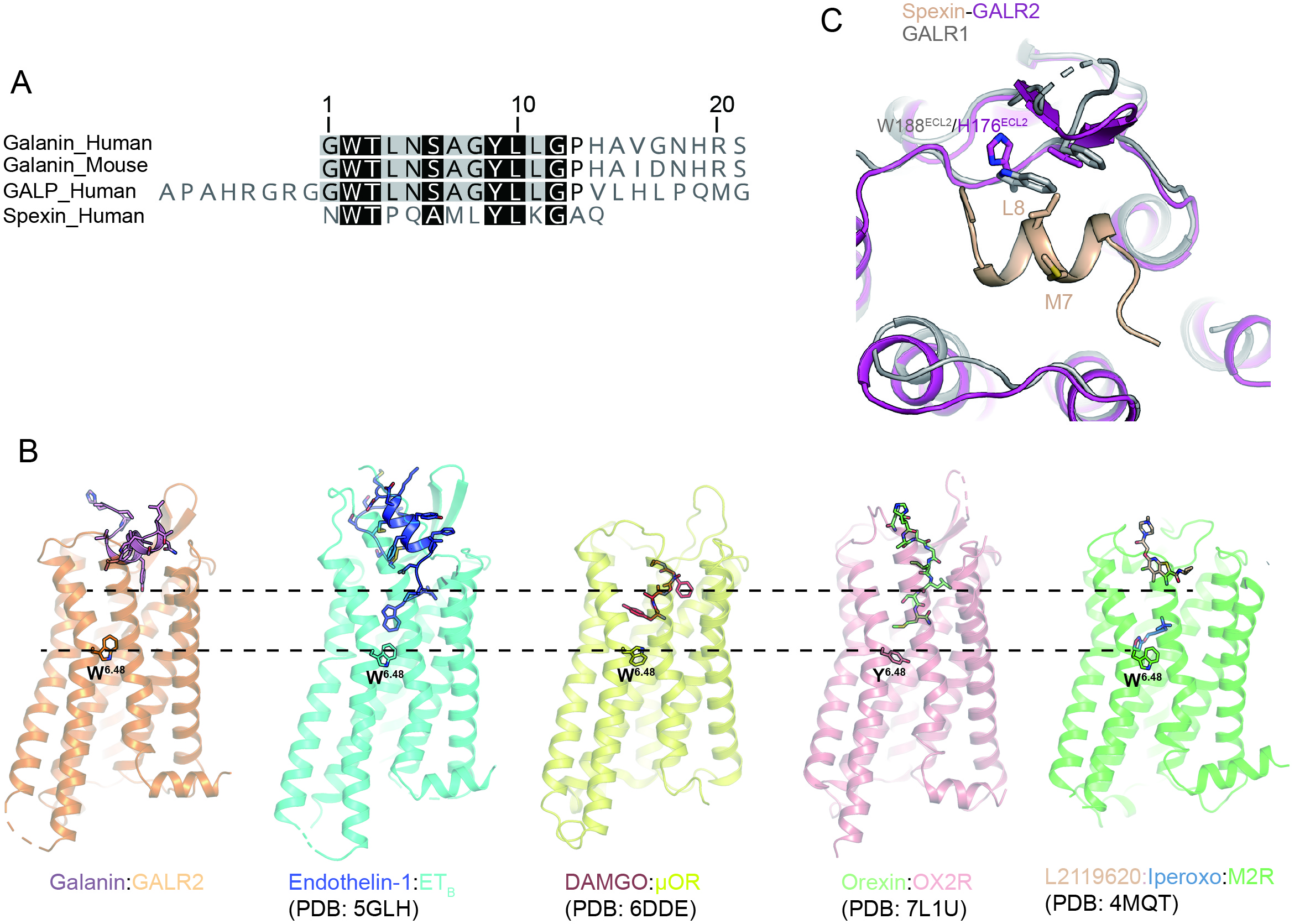

**Fig. S4.** **The unique feature of galanin binding site and binding pose**.

(A) Sequence alignment of the galanin peptide family.

(B) Comparison of the ligand binding among several GPCRs from the class A family. The toggle switch W^6.48^ is shown as sticks.

(C) Structure overlay of spexin-GALR2 and GALR1. L8 in spexin clashes with W188^ECL2^ in GALR1.

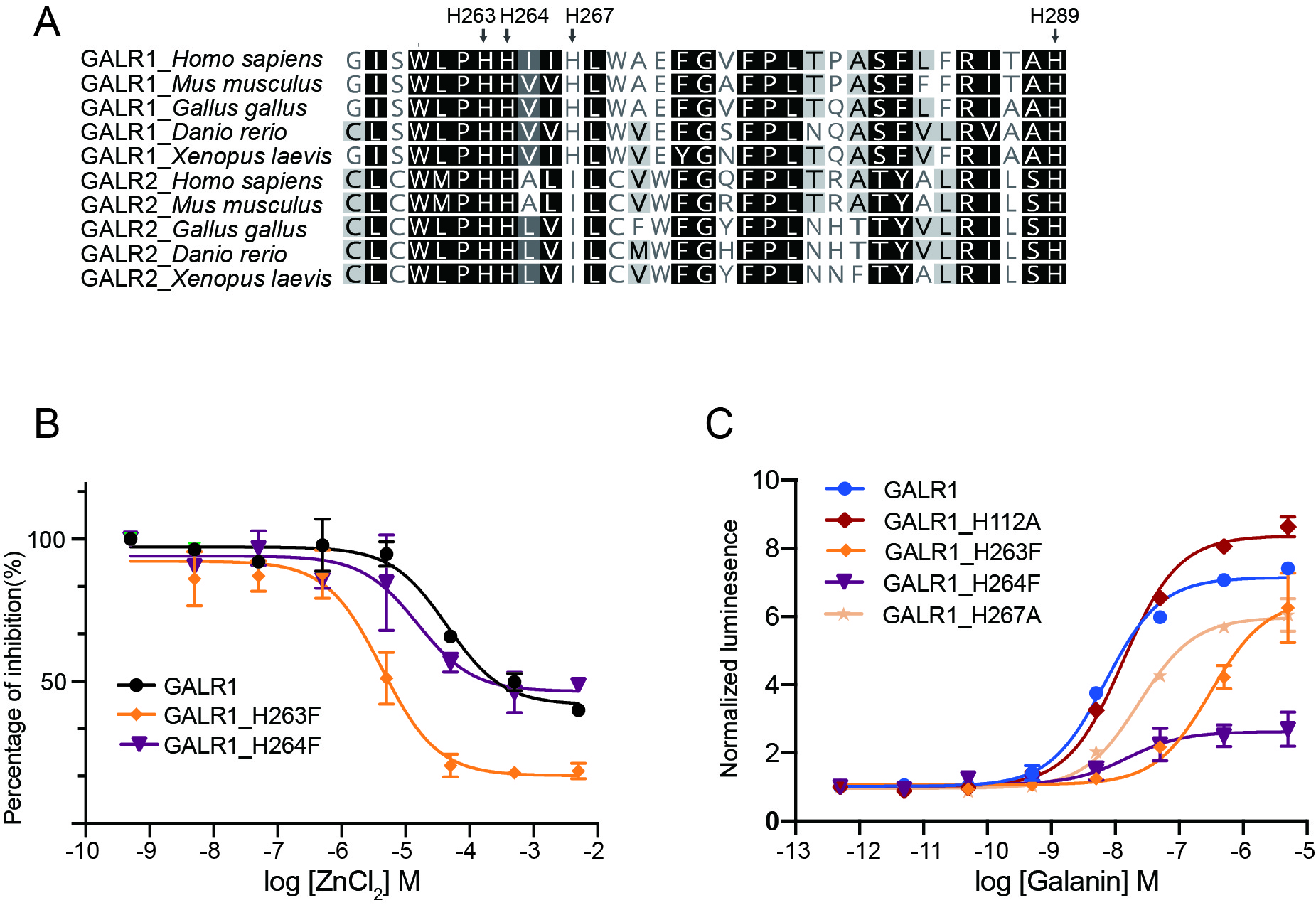

**Fig. S5. Allosteric modulation of GALR1 by zinc.**

(A) Sequence alignment of the histidine-enriched region in GALR1 and GALR2 from multiple species.

(B) The actions of increasing concentration of zinc on activation of GALR1 and its mutants by galanin.

(C) The effects of mutations of histidine residues in GALR1 on receptor activation by galanin.

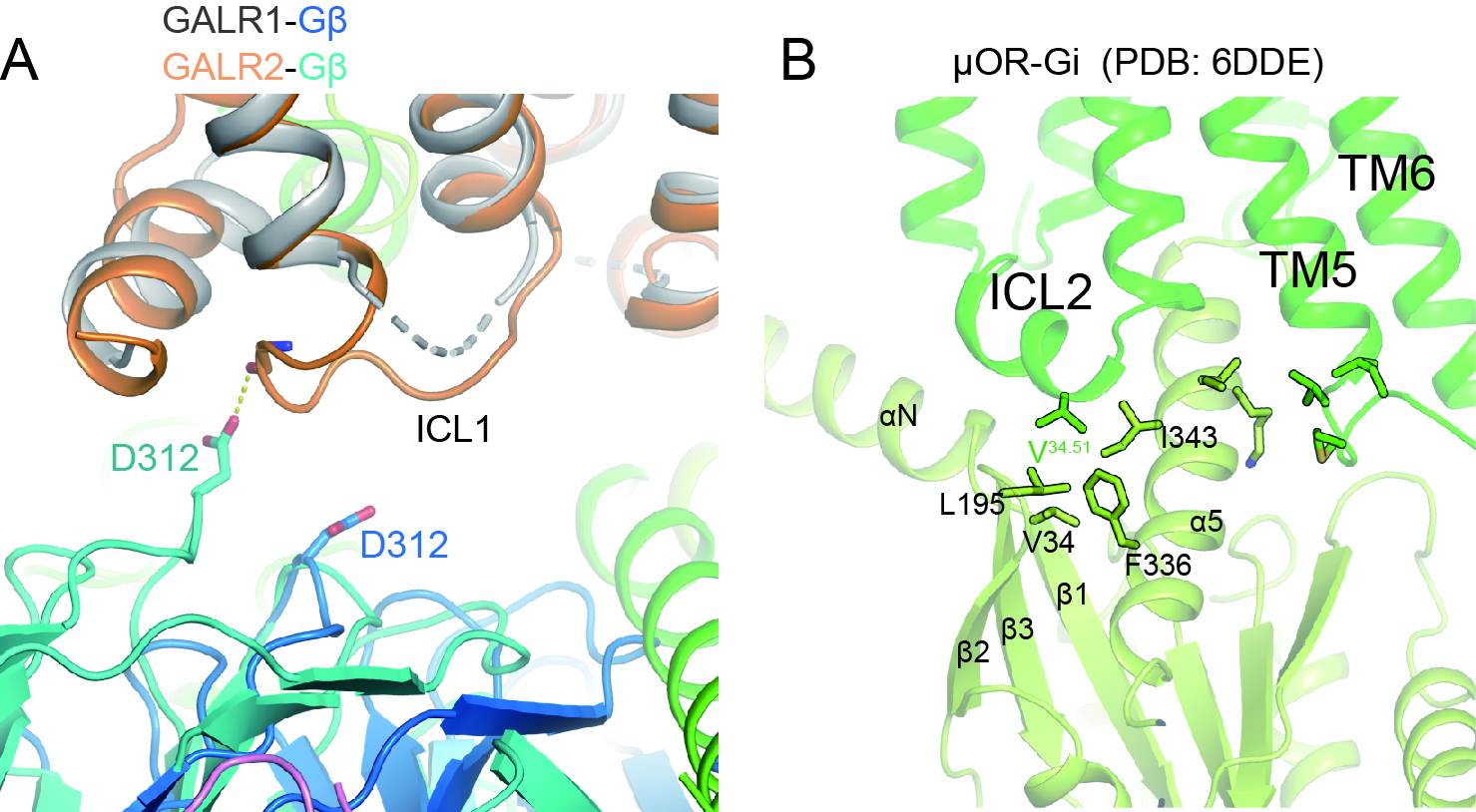

**Fig. S6. Comparison of structures of the GALR1-Go and the GALR2-Gq complexes.**

(A) Close-up view of the ICL1 and Gβ1 interface. Dash line represents hydrogen bond.

(B) Interaction details between ICL2 in μ opioid receptor (μOR) and Gi.

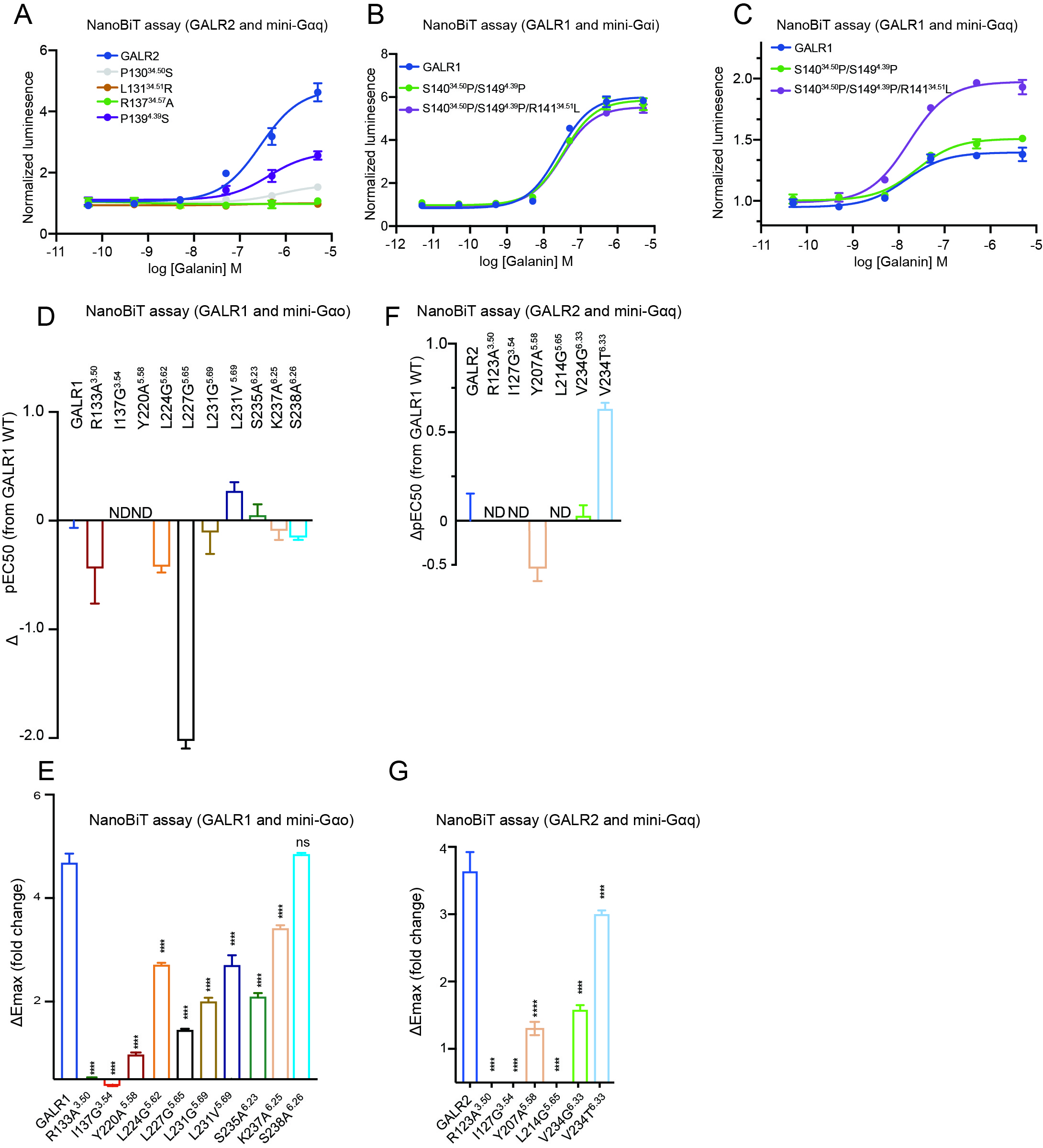

**Fig. S7. The effects of mutations in the GALRs-Gα interface on coupling efficiency**.

(A) Mutations in ICL2 of GALR2 reduce Gq coupling efficiency.

(B) Mutations in ICL2 of GALR1 have little effect on Gi coupling efficiency.

(C) Substitution of ICL2 in GALR1 with that in GALR2 increases coupling efficiency of GALR1-Gq.

(D) and (E) NanoBiT assay for GALR1 and its mutants induced by galanin. The smBit and lgBit are fused to GALR1 and mini-αG, respectively. ΔpEC50 represents differences in galanin potency for each mutant relative to the GALR1 WT. Emax indicates differences in maximum fold change for each mutant relative to the vehicle treatment.

(F) and (G) NanoBiT assay for GALR2 and its mutants.

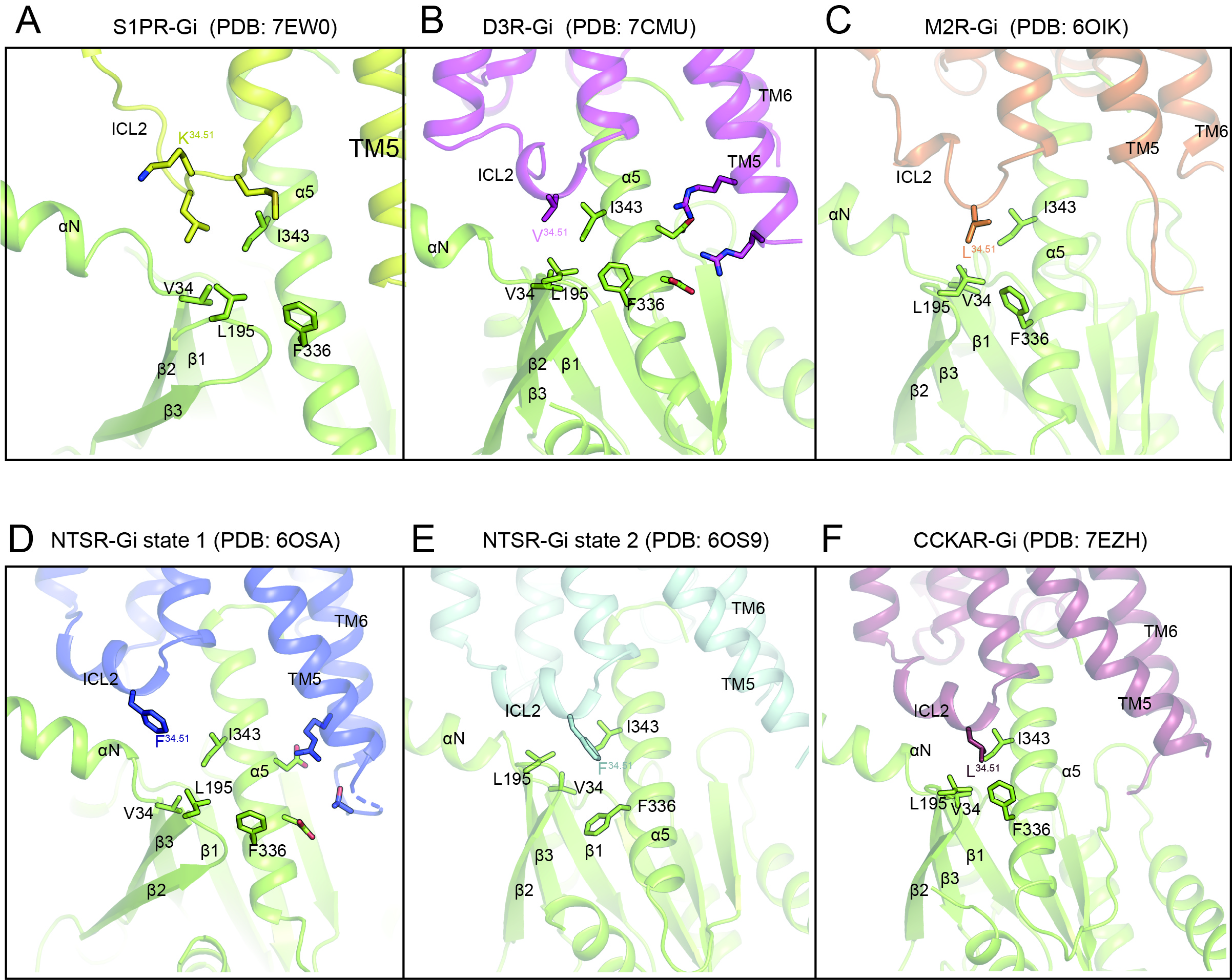

**Fig. S8. Classification of class A Gi-coupled receptors.**

(A) Structure of the S1PR-Gi complex. S1PR belongs to the first class. ICL2 forms a random coiled structure.

(B) and (C), Structures of D3R-Gi and M2R-Gi. D3R and M2R belong to the second class.

(D)-(F) Structures of NTSR-Gi (state 1), NTSR-Gi (state 2) and CCKAR-Gi. NSTR and CCKAR belong to the third class.

**Table S1. Cryo-EM data collection and refinement statistics.**

EM data collection statistics

| Protein | Galanin-GALR1-Go | Galanin-GALR2-Gq | Spexin-GALR2-Gq |
| --- | --- | --- | --- |
| EMDB |  |  |  |
| Microscope | FEI  Titan Krios | FEI Titan Krios | FEI Titan Krios |
| Voltage (kV) | 300 | 300 | 300 |
| Detector | Gatan K3 | Gatan K3 | Gatan K3 |
| Magnification (nominal) | 64000 | 64000 | 64000 |
| Pixel size (Å/pix) | 1.087 | 1.087 | 1.087 |
| Flux (e^-^/pix/sec) | 22 | 22 | 22 |
| Frames per exposure | 32 | 32 | 32 |
| Exposure (e^-^/ Å^2^) | 50 | 50 | 50 |
| Defocus range (μm) | 1.0-2.2 | 1.2-2.3 | 0.7-2.2 |
| Micrographs collected | 2287 | 1337 | 1139 |
| Particles extracted/final | 426,045 | 571,842 | 332,603 |
| Map sharpening B-factor | -171.6 | -176.9 | -200.2 |
| Unmasked resolution  at 0.143 FSC (Å) | 3.4 | 3.4 | 3.7 |
| masked resolution  at 0.143 FSC (Å) | 3.3 | 3.3 | 3.5 |

Model refinement and statistics

|  | Galanin-GALR1-Go | Galanin-GALR2-Gq | Spexin-GALR2-Gq |
| --- | --- | --- | --- |
| PDB |  |  |  |
| Composition |  |  |  |
| Amino acids | 1126 | 1139 | 1137 |
| Ligand | 0 | 0 | 0 |
| RMSD bonds (Å) | 0.003 | 0.003 | 0.004 |
| RMSD angles (º) | 0.691 | 0.633 | 0.968 |
| Mean B-factors |  |  |  |
| Amino acids | 86.88 | 69.83 | 111.48 |
| ligand | 0 | 0 | 0 |
| Ramachandran |  |  |  |
| Favored (%) | 96.29 | 97.67 | 97.4 |
| Allowed (%) | 3.71 | 2.33 | 2.6 |
| Outliers (%) | 0 | 0 | 0 |
| Rotamer Outliers (%) | 0.21 | 0 | 0 |
| Clash score | 11.97 | 11.03 | 16.34 |
| C-beta outliers (%) | 0 | 0 | 0 |
| CC (mask) | 0.72 | 0.76 | 0.76 |
| MolProbity score | 1.84 | 1.63 | 1.83 |
| EMRinger score | 2.19 | 2.75 | 1.69 |
